# The major surface protein of malaria sporozoites is GPI-anchored to the plasma membrane

**DOI:** 10.1101/2024.05.21.595204

**Authors:** Rupa Nagar, Stefano S. Garcia Castillo, Maria Pinzon-Ortiz, Sharon Patray, Alida Coppi, Sachie Kanatani, Robert L. Moritz, Kristian E. Swearingen, Michael A. J. Ferguson, Photini Sinnis

## Abstract

Glycosylphosphatidylinositol (GPI) anchor protein modification in *Plasmodium* species is well known and represents the principal form of glycosylation in these organisms. The structure and biosynthesis of GPI anchors of *Plasmodium* spp. has been primarily studied in the asexual blood stage of *P. falciparum* and is known to contain the typical conserved GPI structure of EtN-P-Man3GlcN-PI. Here, we have investigated the circumsporozoite protein (CSP) for the presence of a GPI-anchor. CSP is the major surface protein of *Plasmodium* sporozoites, the infective stage of the malaria parasite. While it is widely assumed that CSP is a GPI-anchored cell surface protein, compelling biochemical evidence for this supposition is absent. Here, we employed metabolic labeling and mass-spectrometry based approaches to confirm the presence of a GPI anchor in CSP.

Biosynthetic radiolabeling of CSP with [^3^H]-palmitic acid and [^3^H]-ethanolamine, with the former being base-labile and therefore ester-linked, provided strong evidence for the presence of a GPI anchor on CSP, but these data alone were not definitive. To provide further evidence, immunoprecipitated CSP was analyzed for presence of *myo*-inositol (a characteristic component of GPI anchor) using strong acid hydrolysis and GC-MS for a highly sensitive and quantitative detection. The single ion monitoring (SIM) method for GC-MS analysis confirmed the presence of the *myo*-inositol component in CSP. Taken together, these data provide confidence that the long-assumed presence of a GPI anchor on this important parasite protein is correct.

## Introduction

A glycosylphosphatidylinositol (GPI) membrane anchor is a post-translational modification ubiquitous among eukaryotes that provides stable association between proteins and the outer leaflet of the plasma membrane [reviewed in (1)]. Despite this commonality, GPI-anchored proteins are functionally diverse, having roles in signal transduction, immune responses, formation of plasma membrane microdomains, and the pathobiology of parasitic infections. In some protozoan parasites, like African trypanosomes, GPI-anchored surface proteins form a dense surface coat that is hypothesized to protect the parasite from host defenses (2). GPI anchors share a core structure of ethanolamine-phosphate (EtNP) linked to three mannose (Man) residues, a glucosamine residue (GlcN) and a phosphatidylinositol (PI) component (1). This core structure of EtN-P-6Manα1-2Manα1-6Manα1-4GlcNα1-6PI can be elaborated in species- and tissue-specific ways, including additional sugar moieties, variations in lipid component of the PI (diacyl-glycerol, *lyso*-acylglycerol, alkylacyl-glycerol and ceramide) and the presence or absence of fatty acid attached to the *myo*-inositol ring of the PI moiety (1). The latter renders GPI anchors resistant to the action of PI-specific phospholipase C (PI-PLC) and makes these anchors, accordingly, harder to identify. In *Plasmodium* spp., about 30 proteins are anchored to the plasma membrane with GPIs, with most of these identified in the asexual blood stages of the parasite (3). The structure and biosynthesis of *Plasmodium* spp. GPI anchors has mostly been studied in the merozoites of *P. falciparum*. The structure of the GPI anchor of the *P. falciparum* merozoite surface protein-1 and -2 (MSP-1 and MSP-2), and the mature GPI anchor precursor molecule, is EtN-P-6(Manα1-2)Manα1-2Manα1-6Manα1-4GlcNα1-6(acyl)PI where the PI contains diacylglycerol and has a fatty acid attached to the inositol ring (4–7) (**Fig. 2*A***). Similar structural data is also available for the GPI of *P. chabaudi* (8).

The circumsporozoite protein (CSP) is the major surface protein of *Plasmodium* sporozoites, the infective stage of the malaria parasite. Sporozoites, inoculated into the skin of the vertebrate host by an infected mosquito, are motile and must enter blood vessels to go to the liver where they enter hepatocytes and develop into the next life cycle stage (9). Transmission from mosquito to vertebrate host is a bottleneck for the parasite and thus a good target for intervention. Indeed, the only licensed malaria vaccines, RTS,S, and R21, are both subunit vaccines based on CSP that target sporozoites (10, 11). Previous studies have found that CSP is critical for sporozoite development in the mosquito (12) and that controlled exposure of its carboxy terminal adhesion domain functions to guide sporozoites as they migrate from the mosquito midgut to the mammalian liver (13). Though CSP is an abundant surface protein, it does not contain a transmembrane domain and is assumed to be linked to the plasma membrane via a GPI anchor since it possesses a canonical GPI anchor addition sequence at its carboxy terminus (14, 15). Nonetheless, biochemical evidence for the existence of this anchor is absent due to our reliance on harvesting sporozoites from mosquito salivary glands, which limits the amount of parasite material available for analysis.

Given these limitations, genetic approaches were taken to investigate how CSP was anchored to the plasma membrane. Expression of a fusion protein consisting of a portion of human growth factor and the last 28 amino acids of CSP, containing the putative GPI anchor addition sequence, in COS cells resulted in low levels of the fusion protein on the cell surface and metabolic labeling revealed a weak ethanolamine signal (16). Although the majority of the fusion protein was retained intracellularly, the data suggested that the putative CSP-GPI addition sequence could direct the addition of a GPI anchor in a heterologous mammalian expression system. Following this, the putative GPI-addition sequence was mutated in the endogenous *csp* gene, being deleted or replaced by a transmembrane domain (17). In both cases sporozoites failed to develop in oocysts, resembling a *csp* knockout phenotype (12). Though this study showed that the putative GPI-addition sequence is critical for CSP function in sporozoite development, it did not demonstrate the presence of a GPI-anchor. Thus, a full 36 years after the discovery of GPI anchors, it is still not known whether CSP is linked to the plasma membrane using this unique lipid modification. Here, we use metabolic labeling and an ultra-sensitive *myo*-inositol detection methodology to provide evidence that CSP is indeed modified by a GPI anchor.

## Results and Discussion

### Reanalysis of proteomic data for CSP

To begin, we determined whether peptides containing the putative GPI addition sequence of CSP were detected in our published proteomic datasets, with a lack of detection providing support for processing of the CSP’s C-terminus. Typical mass spectrometry-based proteomics workflows digest proteins with trypsin (which cleaves C-terminal to Arg and Lys residues) and the resulting tryptic peptides are separated, detected and fragmented. The peptide mass and fragmentation pattern allow sequence identification. In the many proteomic analyses of *Plasmodium* sporozoites that have been published, CSP is among the most abundant proteins and virtually all of the C-terminal region has been detected by tryptic peptides and semi-tryptic fragments. Notably, however, the C-terminal peptide itself, including the predicted GPI addition (ω) site has never been detected in any experiment in any *Plasmodium* species. The protein sequence coverage of the CSP C-terminus from proteomic analyses of both salivary gland and oocyst sporozoites from human-infective *P. falciparum* (18–20) and *P. vivax* (21) as well as rodent-infective *P. yoelii* (18, 22) and *P. berghei* (23) are shown in (**Fig. 1**). These data include global proteome analyses, as well as proteomes of surface proteins enriched by chemical labeling. In all species, tryptic peptides have been detected that cover the sequence up to and including a conserved Lys residue immediately preceding a conserved Cys residue that is predicted to be the ω-site of processing for attachment of the GPI. However, the predicted ω-site Cys residue and the 23 amino acids downstream of it, which comprises a single predicted tryptic peptide in all four species, have never been detected suggesting that the C-terminus of CSP has been processed in sporozoites rendering this predicted tryptic peptide undetectable. Mining of sporozoite transcriptome and proteome databases (18, 20 24, 25) demonstrate that the entire GPI anchor precursor biosynthesis machinery, and GPI transamidase, are expressed in sporozoites (**Table S1**; data from https://plasmodb.org (26)). Although the absence of evidence for a detectable C-terminal tryptic peptide cannot be taken as a proof, these lines of evidence are consistent with CSP being a GPI-anchored protein, most likely with the GPI attached to the α-carboxyl group of the Cys residue.

**Figure 1.**
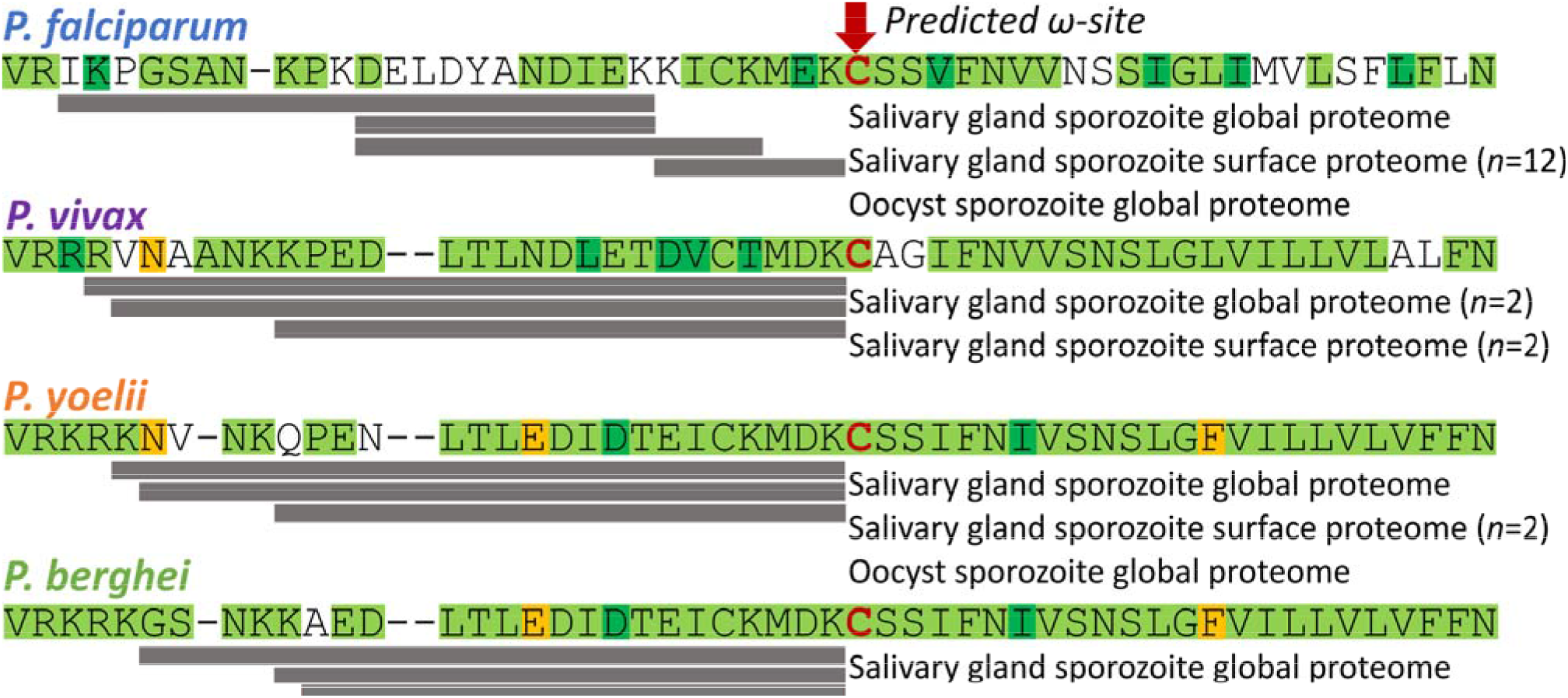
Proteomic evidence for processing of the CSP C-terminus at the putative GPI anchor addition sequence. The sequences of the C-termini of CSP from different *Plasmodium* species are shown, beginning at the basic residues after the type I thrombospondin repeat (TSR). Published mass spectrometry-based proteomic analyses of sporozoites failed to detect peptides corresponding to the predicted GPI addition site (red arrow) and downstream residues, consistent with prior processing of the protein and removal of the putative GPI-addition sequences. Grey bars indicate detected tryptic peptides (cleavage after Lys and Arg) identified across the indicated datasets, which are listed under the putative GPI-addition sequence for each species. Though not all tryptic peptides were detected in every dataset, the grey bars show a consensus of the data from species for which multiple proteomes are available. Proteomes for which replicates are not listed are n=1. Amino acid sequences are color coded to indicate degree of conservation across species: Green = conserved residues, Dark green = substitutions with similar residues, Yellow = non-similar residues that are conserved between two of the four species, No color = not conserved.

**Figure 2.**
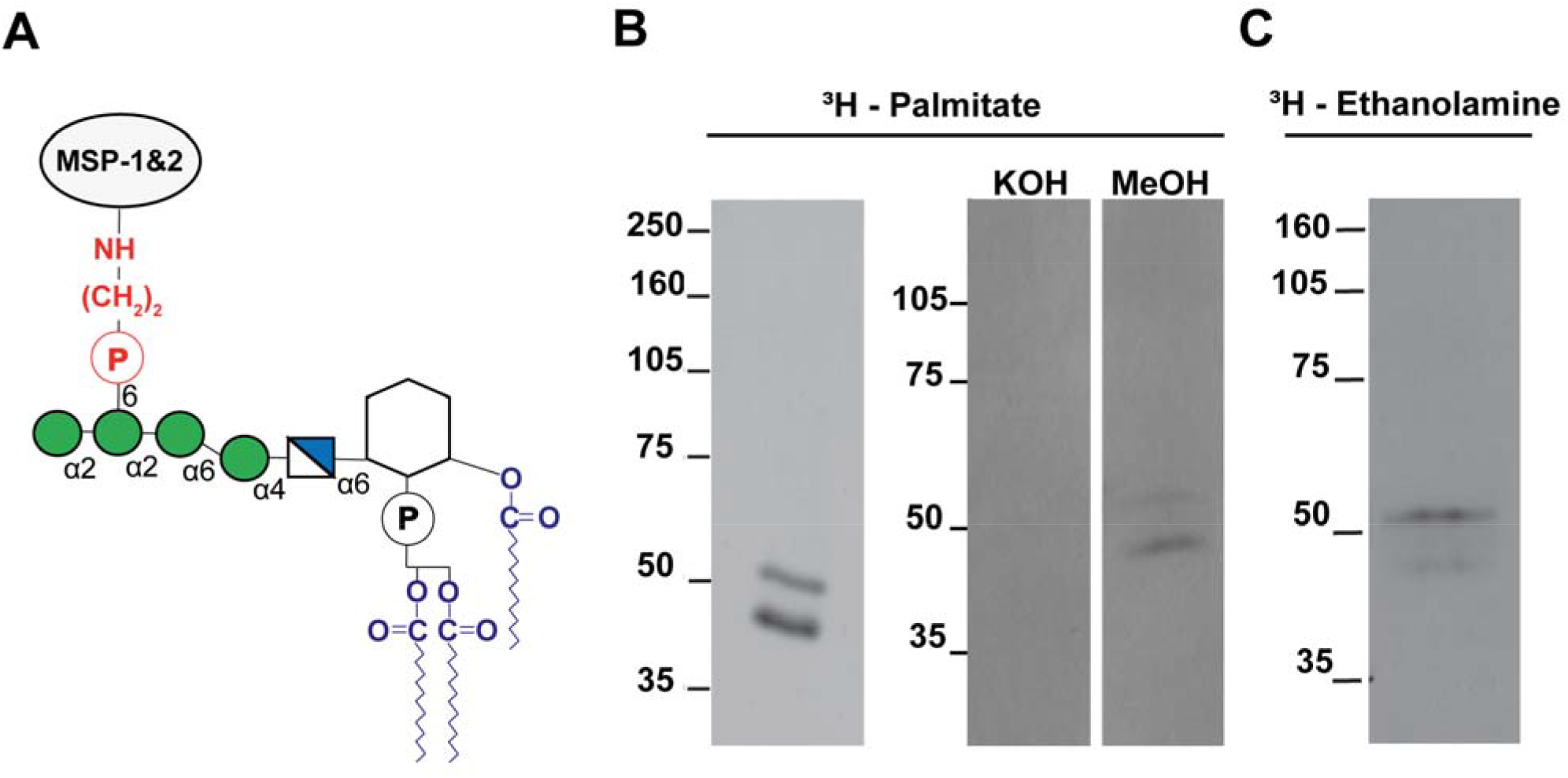
Metabolic labeling with [^3^H]-palmitic acid and [^3^H]-ethanolamine. (**A**) Cartoon of a GPI anchor, based on the structure for the GPI anchor of *P. falciparum* merozoite major surface proteins 1 and 2 (1). The portions predicted to be labeled by [^3^H]-palmitate and [^3^H]-ethanolamine are shown in blue and red, respectively. The ethanolamine phosphate bridge (shown in red) links the carboxy terminus of the processed protein to a phosphatidylinositol group via a carbohydrate chain that contains mannose (Man, green circles) and glucosamine (GlcN, blue and white square). The three fatty acids (shown in blue) within the phosphatidylinositol group, two attached to glycerol and one to the *myo*-inositol ring, anchor the protein to the cell membrane. (**B**) [^3^H]-palmitate labeling and base-hydrolysis of labeled CSP. *P. berghei* sporozoites were metabolically labeled with [^3^H]-palmitate coupled to fatty-acid free BSA for 12 h, washed, lysed, and CSP was immunoprecipitated. Left panel: the immunoprecipitate was analyzed by SDS-PAGE followed by fluorography. Right panel: the immunoprecipitate was equally divided into 2 lanes and separated by SDS-PAGE. The gel was fixed, cut in half and one half was incubated in 0.2M KOH in methanol and the other in methanol alone. The gel was then washed, soaked with Amplify™ and dried for fluorography. (**C**) [^3^H]-ethanolamine labeling. *P. berghei* sporozoites were metabolically labeled with [^3^H]-ethanolamine for 12 h followed by immunoprecipitation of CSP and analysis by SDS-PAGE and fluorography.

### [^3^H]-palmitic acid and [^3^H]-ethanolamine are incorporated into CSP

We performed metabolic labeling experiments to assess incorporation of radiolabeled GPI precursors into CSP. Using the rodent malaria parasite *Plasmodium berghei*, we harvested sporozoites from the salivary glands of infected mosquitoes and incubated them with the indicated radiolabeled precursor. Following this, CSP was immunoprecipitated and analyzed by SDS-PAGE and fluorography. CSP runs as 2 to 3 bands by SDS-PAGE (27, 28), with the lower band being the product of a proteolytic cleavage event that results in the removal of 10 kDa from the N-terminus (29). Since CSP cleavage occurs after the protein has been trafficked to the parasite surface (13, 29), one would expect both bands to be labeled with the GPI precursors. As shown in **Fig. 2*B*** (left panel), both CSP bands are labeled with [^3^H]-palmitate, predicted to be incorporated into the GPI lipid moiety that anchors the protein to the outer leaflet of the plasma membrane.

To determine whether the [^3^H]-palmitate was covalently linked to CSP by an ester linkage, we performed base-hydrolysis of [^3^H]-palmitate-labeled CSP. CSP was immunoprecipitated from metabolically-labeled sporozoites and the immunoprecipitate was loaded onto two lanes of an SDS-PAGE gel. After electrophoresis, the gel was cut into separate lanes, and one lane was subjected to base hydrolysis using KOH in methanol and the other control lane was incubated with methanol alone. As shown in **Fig. 2*B*** (right panel), base hydrolysis removes the [^3^H]-palmitate label, consistent with its being linked via an oxy-ester or thio-ester bond.

While the presence of ester-linked [^3^H]-palmitate is consistent with CSP being anchored to the parasite surface by a GPI anchor, it does not constitute definitive proof. Palmitic acid can also directly modify serine, threonine, or cysteine residues via ester bonds. Though direct palmitoylation of serine, threonine, or cysteine residues is canonically found on transmembrane proteins or proteins associated with the inner leaflet of the plasma membrane, neither of which is likely the case for CSP, we cannot formally exclude the possibility that the [^3^H]-palmitate labeling of CSP is not a result of direct palmitoylation of one or more of these amino acids. We therefore also attempted to biosynthetically label CSP with [^3^H]-ethanolamine, a core component of GPI anchors (**Fig. 2*A***). *P. berghei* sporozoites were incubated with [^3^H]-ethanolamine and processed as outlined above. As shown in **Fig. 2*C***, [^3^H]-ethanolamine is incorporated into CSP, consistent with being GPI-anchored to the sporozoite plasma membrane.

### Chemical detection of myo-inositol in CSP

While the incorporation of [^3^H]-ethanolamine into CSP is highly suggestive of it being GPI anchored, [^3^H]-ethanolamine can be incorporated into proteins via a different mechanism (30, 31) albeit this occurs on intracellular proteins such as elongation factor-1α rather than plasma membrane proteins. For this reason, we sought to confirm the presence of a definitive GPI anchor component (*myo*-inositol) in CSP by direct chemical means.

For this experiment we used *Plasmodium falciparum* sporozoites because of the higher parasite numbers we obtain in mosquitoes. Approximately 500 mosquitoes were dissected and CSP was immunoprecipitated from harvested sporozoites. A sample of immunoprecipitated CSP was taken for SDS-PAGE and blotted onto polyvinylidene diflouride (PVDF) membrane. Five sections of the blot, together with equivalent sections from a lane with no CSP sample, were excised (**Fig. 3*A***). These PVDF strips were transferred to reaction vials and, after the addition of an internal standard of [1,2,3,4,5,6-^2^H]-*myo*-inositol (D_6_-*myo-*inositol), individually subjected to strong acid hydrolysis (6N HCl, 110 °C, 24 h) to release *myo*-inositol from any molecules on the PVDF membrane as free *myo*-inositol. The hydrolysates were removed from the PVDF strips, dried and derivatized with trimethylsilyl (TMS) reagent prior to analysis by gas chromatography-mass spectrometry (GC-MS) in selected ion monitoring (SIM) mode to quantitate free *myo*-inositol released from the PVDF membrane strips, (**Fig. 3*B***). The ions monitored were at *m/z* 305 and 318, both prominent electron impact fragment ions of unlabeled *myo*-inositol-TMS_6_, and *m/z* 307 and 321 which are the corresponding ions from the D_6_-*myo*-inositol-TMS_6_ internal standard (**Fig. 3*C***). The deuterated *myo*-inositol-TMS_6_ derivative elutes slightly earlier than the non-deutero equivalent.

**Figure 3.**
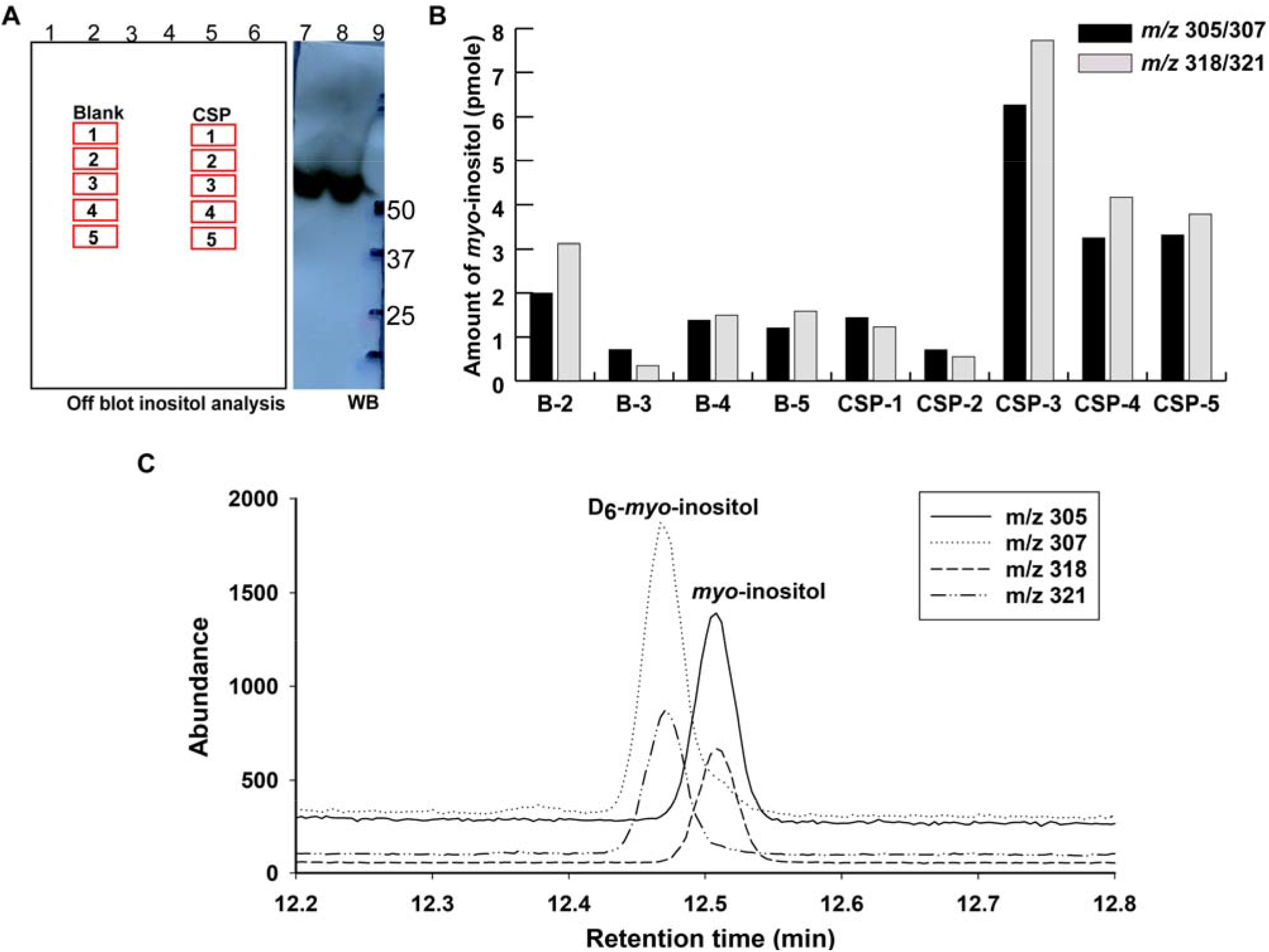
*Myo*-inositol analysis of immunoprecipitated CSP. (**A**) A representation of the PVDF membrane carrying the immunoprecipitated CSP and showing the positions of the strips used for *myo*-inositol analysis. Lanes 1 and 9, molecular weight markers; lanes 2-4 and 6, blank; lane 5, immunoprecipitated CSP equivalent to 3.9x10^7^ sporozoites; lanes 7 and 8, immunoprecipitated CSP equivalent to 1x10^4^ and 5x10^4^ *P. falciparum* sporozoites, respectively. Lanes 7-9 were subjected to Western blotting and were used to estimate the position of CSP on the blot. (**B**) Amounts of *myo*-inositol present on the strips of PVDF blot for the immunoprecipitated CSP sample and negative control (blank lane) were measured using a selected ion monitoring (SIM) GC-MS approach. (**C**) A representative extracted ion chromatogram (EIC) of the CSP-3 sample, where *m/z* 305 and *m/z* 318 represent the native H_6_-*myo*-inositol of the CSP-3 sample and *m/z* 307 and 321 represent the D_6_-*myo*-inositol internal standard.

The results of this analysis showed a sharp increase in *myo*-inositol content in strip 3 of the immunoprecipitated CSP sample lane, which coincided with the position of CSP by the Western blot with anti-CSP antibody run in parallel (**Fig. 3*A***, lanes 7 and 8). The level of *myo*-inositol content in strips 4 and 5 were lower than for strip 3 but remained higher than for strips 1 and 2 (**Fig. 3*B*)**. We think this may be due to small amounts of proteolytically processed CSP that would also contain the GPI anchor. The presence of *myo*-inositol in the CSP material detected here, together with the ability to label CSP with [^3^H]-palmitate and [^3^H]-ethanolamine, provide strong evidence for the presence of a GPI anchor on the CSP protein. Taking the data from strip 3 of 6 pmol of *myo*-inositol above background from 3.9 x 10^7^ sporozoites (**Fig. 3*B***), we can calculate a minimum copy number of GPI anchors, and thus of CSP itself, of about 100,000 copies per cell. However, this is an absolute minimum estimate because it assumes 100% yield throughout all stages of the processing and analysis, which is very unlikely given the lengthy procedure involving several washes after immunoprecipitation, SDS-PAGE and transfer to PVDF.

### Inhibitors of GPI synthesis do not impact sporozoite motility

Sporozoites move by a substrate-based gliding motility that is required for exit from the inoculation site and entry into hepatocytes (reviewed in (32)). As they move, CSP is shed from the sporozoite surface leaving trails (33), leading us to hypothesize that inhibitors of GPI anchor synthesis might reduce sporozoite motility. We tested salicylic hydroxamic acid (SHAM) an inhibitor of N-acetylglucosaminyl-phosphatidylinositol de-N-acetylase, and manogepix, gepinacin and MMV892853, inhibitors of inositol acyltransferase (34–37) in motility assays with *P. berghei* and *P. falciparum* sporozoites. However, we did not observe inhibition of motility in the presence of up to 100 μM of each inhibitor (**Fig. S1**). Without direct evidence that these conditions reduced or prevented GPI addition to CSP, these data are difficult to interpret. Nevertheless, we present them here for consideration.

In conclusion, the presence of *myo*-inositol in immunoprecipitated and SDS-PAGE isolated CSP material, together with the ability to label CSP with [^3^H]-ethanolamine and [^3^H]-palmitate, provides compelling data that CSP is GPI anchored. While this is not unexpected given the presence of a putative GPI anchor addition signal peptide sequence at its C-terminus, we now provide the necessary biochemical evidence to support this long-standing assumption.

## Experimental procedures

### Mosquito infections with Plasmodium berghei

*Anopheles stephensi* (Liston strain) mosquitoes were reared in the insectary at the Johns Hopkins Malaria Research Institute and infected with *Plasmodium berghei* ANKA strain as described previously (9). Briefly, female 5 to 6 week old Swiss Webster mice (Taconic) with gametocytemias of 0.1 to 0.3% were fed upon by *Anopheles stephensi* mosquitoes which were maintained on a 10% sugar solution at 20°C and 80% humidity with a 14:10 hr light:dark cycle including a 2 hr dawn/dusk transition. On days 18 to 22 post blood meal sporozoites were harvested from mosquito salivary glands for use in experiments. Animal work was approved by the Johns Hopkins University Animal Care and Use Committee (protocols #M011H467 and #M014H363), which is fully accredited by the Association for the Assessment and Accreditation of Laboratory Animal Care.

### Mosquito infections with Plasmodium falciparum

Gametocyte cultures and mosquito infections were performed as previously described (38), maintaining *P. falciparum* NF54 blood stage cultures in 4% human erythrocytes in RPMI-1640 culture medium (2 mM L-glutamine, 50 mg/L hypoxanthine, 25 mM HEPES, 0.225% NaHCO_3_, and 10% *v/v* human serum) at 37°C in a candle jar made of glass desiccators. Gametocyte cultures were initiated at 0.5% asexual parasitemia from low passage stock and maintained up to day 18 with daily media changes but without any addition of fresh erythrocytes. Day 15 to 18 cultures, containing largely mature gametocytes, were used for mosquito feeds: Cultures were centrifuged at 108 x *g* for 4 min and the parasite pellet was resuspended to final gametocytemia of 0.3% in a mixture of human O^+^ RBCs supplemented with 50% *v/v* human serum. Gametocytes were fed to *Anopheles stephensi* mosquitoes that had been starved overnight, using glass membrane feeders. Unfed mosquitoes were removed after feeding and mosquitoes were maintained as outlined above, except that the incubator temperature was 25°C. Human erythrocytes used to set up the cultures are collected weekly from healthy donors under an institutional review board–approved protocol.

### Radiolabeling of CSP

Salivary glands of *P. berghei* infected mosquitoes were dissected on days 18 to 22 post blood meal, homogenized and sporozoites were counted on a hemocytometer. Between 2.5 to 5 x 10^6^ sporozoites were used for each experiment. Sporozoites were metabolically labeled in RPMI-1640 containing 5x Pen/Strep, 25 mM Hepes, and 1% BSA pH 7.4 in a total volume of 500 μL with 100 μCi per ml of [^3^H]-palmitate or [^3^H]-ethanolamine. Prior to addition of [^3^H]-palmitate, it was coupled to defatted BSA (Sigma A8806) by drying (under N_2_), dissolving in 10 μL absolute ethanol and vortexing with the BSA. Sporozoites were incubated in labeling medium overnight (∼16 hrs) at 25°C.

### Immunoprecipitation and SDS-PAGE analysis

Antibodies used for immunoprecipitation were mAb 3D11, specific for the repeat region of *P. berghei* CSP and mAb 2A10, specific for the repeat region of *P. falciparum* CSP. MAbs 3D11 and 2A10 were conjugated to Sepharose 4B using the protocol outlined in (39). Metabolically-labeled sporozoites were washed 3 times in RPMI to remove unincorporated label and resuspended in 400 μL of lysis buffer (1% Triton X-100, 50 mM Tris-HCl pH 8.0) with 150 mM NaCl and a protease inhibitor cocktail (Complete Mini-Tablets, Roche) for 1 hr at 4°C. The lysate was centrifuged for 30 min at 14,000 rpm at 4°C, and the supernatants were incubated with mAb 3D11 or mAb 2A10 conjugated to agarose overnight at 4°C with agitation, and the beads were then washed sequentially with lysis buffer containing 150 mM NaCl, high salt buffer (500 mM NaCl in lysis buffer), lysis buffer without added NaCl, and pre-elution buffer (0.5% Triton X-100, 10 mM Tris-HCl pH 6.8). CSP was eluted with 1% SDS in 0.1 M glycine pH 1.8, neutralized with 1.5 M Tris-HCl pH 8.8, and run on a 7.5% SDS-polyacrylamide gel under nonreducing conditions. For experiments with *P. falciparum*, a 10% SDS-polyacrylamide gel was used. Gels were fixed in 25% methanol/12% acetic acid, enhanced with Amplify (Amersham Pharmacia) for 30 min, dried and exposed to film at minus 80°C for 2 to 3 months.

### Base hydrolysis of [^3^H]-palmitate labeled CSP

Sporozoites were labeled with [^3^H]-palmitate and CSP was immunoprecipitated and run on an SDS-polyacrylamide gel as outlined above. The gel was fixed for 90 min in 25% methanol/12% acetic acid, washed 3 times with deionized H_2_O to remove the acetic acid and cut in half, with each containing half of the immunoprecipitated, eluted CSP. One half was incubated in 0.2 M KOH in methanol and the other in methanol alone, overnight at room temperature. The gel slices were washed 3 times (10 min per wash) with deionized H_2_O, enhanced with Amplify for 30 min, dried and exposed to film as above.

### Myo-inositol analysis

The salivary glands of ∼500 *P. falciparum* infected mosquitoes were lysed using 1% Triton X-100 to yield 4 x 10^7^ sporozoites. CSP was immunoprecipitated with mAb 2A10 conjugated to Sepharose and eluted with 0.1 M glycine pH 1.5 in 1% SDS from the beads. The eluted proteins were separated by NuPAGE bis-Tris 4–12% gradient acrylamide gels (Invitrogen) and transferred to PVDF membrane (Invitrogen). For *myo*-inositol analysis, approximately 3.9 x 10^7^ equivalent sporozoites were loaded in a single lane. Smaller amounts of 1 x 10^4^ and 5 x 10^4^ equivalent sporozoites were loaded in parallel lanes for immunoblotting. After the transfer, the blot was cut in half to separate the lanes for CSP inositol analysis and immunoblotting. The immunoblot was developed using mAb 2A10. The remaining blot containing 3.9 x 10^7^ equivalent sporozoites was subjected to inositol analysis as previously described in (40, 41). Briefly, the estimated CSP band area (as correlated with parallel immunoblot) and equivalent areas from above and below the protein band were cut out. A parallel blank lane was cut in identical manner to use as negative controls for the experiment. Each PVDF strip was washed in methanol and placed in 2 ml glass reactivial containing 250 μL of 6 N HCl and 10 pmoles of D_6_-*myo*-inositol ([1,2,3,4,5,6-^2^H]-*myo*-Inositol from QMX, Labs, Thaxted, UK) internal standard. 10 pmoles of purified sVSG (native H_6_-*myo*-inositol) mixed with 10 pmoles of D_6_-*myo*-inositol internal standard was processed alongside as a standard. The vials were sealed with a Teflon lined cap and hydrolyzed for 24 hr at 110 °C. The hydrolysates were dried in a Speed vac, followed by redrying with 100 μL water and then with 100 μL methanol. The samples were then derivatized with TMS reagent and analyzed using single ion monitoring (SIM) approach on GC-MS (Agilent Technologies, 7890B Gas Chromatography system with 5977A MSD) equipped with Agilent J&W HP-5ms GC Column (30 m x 0.25 mm, 0.25 μm) with He carrier gas at 0.5 mL/min. The native H_6_-*myo*-inositol was detected by the characteristic ions at *m/z* 305 and *m/z* 318 and the D_6_-*myo*-inositol internal standard was detected using *m/z* 307 and *m/z* 321. The amount of H_6_-*myo*-inositol was calculated as follows: (area of *m/z* 305 peak / area of *m/z* 307 peak) x amount of D_6_-*myo*-inositol internal standard; and (area of *m/z* 318 peak / area of *m/z* 321 peak) x amount of D_6_-*myo*-inositol internal standard. The 10 pmol sVSG standard yielded 9.1 and 11.0 pmoles of H_6_-*myo*-inositol, respectively, by this method, providing experimental validation.

### Gliding motility assay

Coverslips were coated overnight with 5 μg/mL of mAb 3D11 [*P. berghei* (42)] or 3 μg/mL mAb 2A10 [*P. falciparum* (43)] in PBS to enrich for capture of shed CSP. On the day of the experiment, mosquito salivary glands were dissected, homogenized to release sporozoites, counted on a hemocytometer and preincubated in HBSS with 2% (wt/vol) BSA at pH 7.4 (untreated control) or 1 μM cytochalasin D or 100 μM of the indicated GPI anchor biosynthesis inhibitor [Salicylic hydroxamic acid (SHAM), MMV892853, Gepinacin and Manogepix] for 30 min at 20°C. Coated coverslips were washed twice with PBS and 65,000 (*P. berghei*) or 75,000 (*P. falciparum*) sporozoites in the continued presence of the inhibitor were added to the coverslips (in a 24-well plate), centrifuged for 5 min at 150 x g and incubated for 1 hr at 37°C. Following this, coverslips were fixed with 4% paraformaldehyde and stained for CSP using biotinylated mAb 3D11 or mAb 2A10 followed by Alexa Fluor 488 streptavidin. Total fluorescence intensity of sporozoites and trails from 25 fields per well were quantified using a fluorescence microscope (Nikon E600) with a 40x objective and ImageJ. Percent motility was calculated by dividing each sample’s fluorescence value by the mean of the untreated control’s fluorescence value. The data were pooled from 3 (*P. berghei*) or 2 (*P. falciparum*) independent experiments.

## Abbreviations

The abbreviations used are

GPI: glycosylphosphatidylinositol
CSP: circumsporozoite protein
MSP: merozoite surface protein
EtN-P: ethanolamine-phosphate
GlcN: glucosamine
Man: mannose
Gal: galactose
PI: phosphatidylinositol
GC-MS: gas chromatography-mass spectrometry
PVDF: polyvinylidene difluoride
TMS: trimethylsilyl
LC-MS/MS: liquid chromatography-tandem mass spectrometry
mAb: monoclonal antibody
sVSG: soluble variant surface glycoprotein

## Data availability

All data pertinent to this work are contained within this article and are available upon request from MAJF and PS.

## Supporting information

This article contains supporting information.

## Acknowledgements

We are grateful to the insectary and parasitology core facilities at the Johns Hopkins Malaria Research Institute and would like to thank Abhai Tripathi, Godfree Mlambo and Chris Kizito for their outstanding work and Bloomberg Philanthropies for supporting these facilities. PS would like to thank the GPI community for helpful discussions along the way, including Drs. Jayne Raper, Jay Bangs, Anant Menon, Alvaro Acosta-Serrano, Martin Low, Clement Bordier, Paul Englund, and Maria Lucia Cardoso de Almeida.

## Author contributions

PS and MAJF conception and design; RN, SSGC, MPO, SP, AC, SK, KES, MAJF and PS data acquisition; RN, SSGC, KES, MAJF and PS data analysis and interpretation; RN, SOGC, MPO, MAJF and PS methodology; RN, MAJF and PS writing–original draft; RLM, MAJF. and PS funding acquisition; RN, MAJF and PS writing–review and editing.

## Funding and additional information

Research reported in this publication was supported by the Bill and Melinda Gates Foundation under award number OPP1139130 (PS and KES), the National Institutes of Health R01 AI 56840 (PS), R01 AI 148489 (KES) and Diversity Supplement to R01 AI132359 (SP), a Johns Hopkins Malaria Research Institute postdoctoral fellowship (SK) and Wellcome Trust Investigator Award 101842/Z13/Z (MAJF).

## Notes

### Competing Interest Statement

The authors have declared no competing interest.

