## Supporting information for "The major surface protein of malaria sporozoites is GPI-anchored to the plasma membrane"

**Table S1:** Mining of transcriptome and proteome databases (https://plasmodb.org) for presence of GPI anchor biosynthesis machinery and GPI transamidase.

| Gene Name | Oocyst Spz Transcriptome | Salivary Gland Spz Transcriptome | Salivary Gland Spz Proteome | Blood Stage Transcriptome | Blood Stage Proteome |
|---|---|---|---|---|---|
| **PF3D7_1032400 PIGA** phosphatidylinositol N-acetyl glucosaminyltransferase subunit A | Yes | Yes | No | Yes | No |
| **PF3D7_1141400** phosphatidylinositol N-acetyl glucosaminyltransferase subunit H | Yes | No | No | Yes | Yes |
| **PF3D7_0618900** phosphatidylinositol N-acetyl glucosaminyltransferase subunit GPI1 | Yes | Yes | No | Yes | Yes |
| **PF3D7_0911000** phosphatidylinositol N-acetyl glucosaminyltransferase subunit C | Yes | Yes | No | Yes | No |
| **PF3D7_0935300** phosphatidylinositol N-acetyl glucosaminyltransferase subunit P | Yes | Yes | No | Yes | No |
| **PF3D7_0624700** N-acetylglucosaminyl-phosphatidylinositol de-N-acetylase | Yes | Yes | No | Yes | Yes |
| **PF3D7_0615300** GPI-anchored wall transfer protein 1, (inositol acyltransferase) | Yes | Yes | No | Yes | No |
| **PF3D7_1210900** GPI mannosyltransferase 1 | Yes | Yes | No | Yes | No |
| **PF3D7_1247300** GPI mannosyltransferase 2 | Yes | Yes | No | Yes | No |
| **PF3D7_1341600** GPI mannosyltransferase 3 | Yes | Yes | No | Yes | No |
| **PF3D7_1214100** GPI ethanolamine phosphate transferase 3 | Yes | Yes | Yes | Yes | Yes |
| **PF3D7_1122100** GPI transamidase component GPI16 | Yes | No | Yes | Yes | Yes |
| **PF3D7_1330700** GPI transamidase subunit PIG-U | Yes | Yes | No | Yes | Yes |

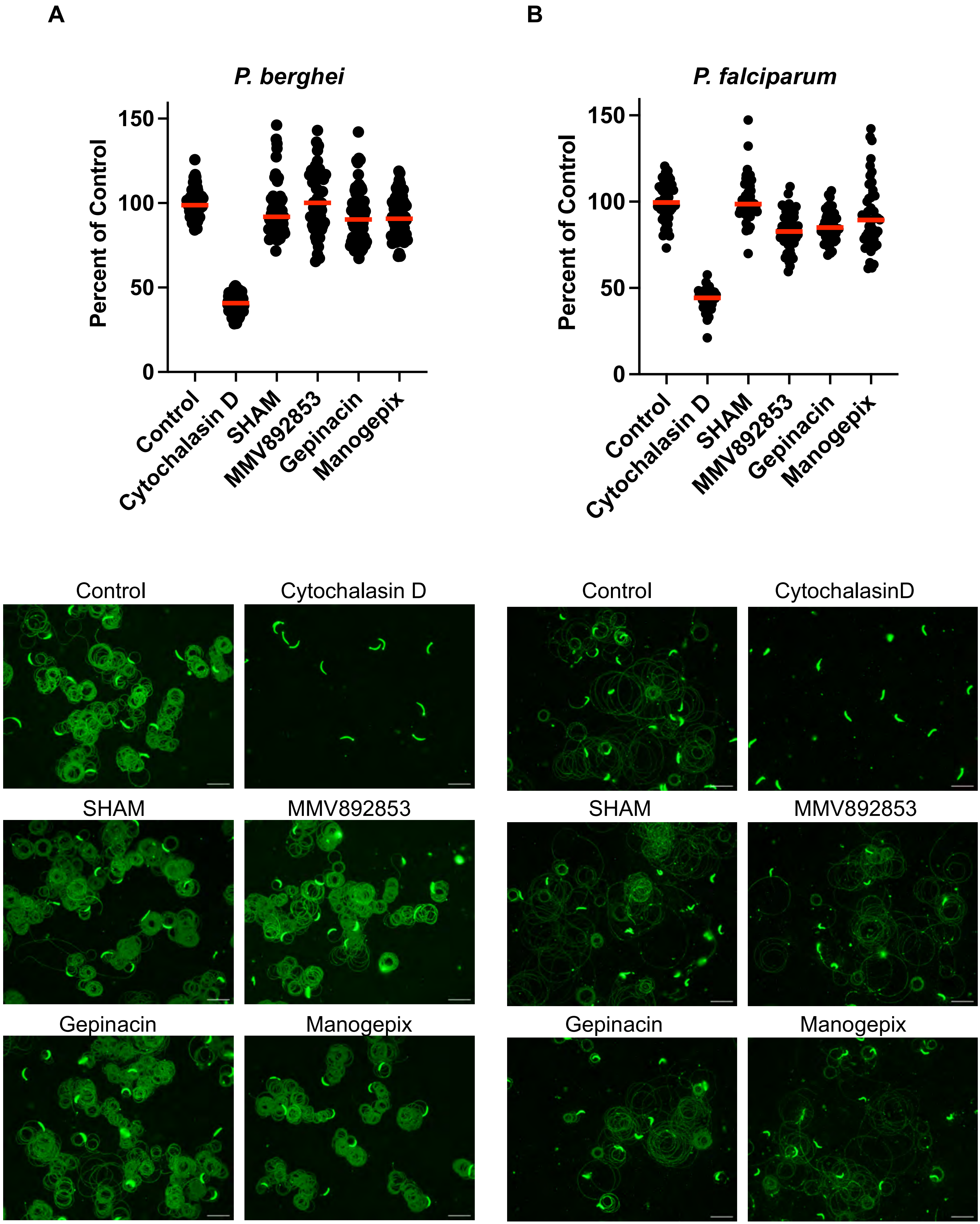
A
B
P. berghei
P. falciparum
Percent of Control
150
100
50
0
Control
Cytochalasin D
SHAM
MMV892853
Gepinacin
Manogepix
Control
Cytochalasin D
SHAM
MMV892853
Gepinacin
Manogepix
Control
CytochalasinD
SHAM
MMV892853
Gepinacin
Manogepix

**Supplementary Figure 1. GPI anchor biosynthesis inhibitors do not inhibit *Plasmodium* sporozoite motility.** *P. berghei* (A) or *P. falciparum* (B) sporozoites were harvested from mosquito salivary glands and incubated with 100 µM of the indicated inhibitor for 30 minutes prior to being added to coated coverslips and allowed to glide for 1 hr at 37°C. Following this coverslips were fixed and stained to visualize sporozoites and the trails they left behind. Shown in the upper panels is total fluorescence intensity from 25 images per well was quantified using Image J. Data were pooled from 3 (*P. berghei*) or 2 (*P. falciparum*) independent experiments and normalized to percent motility by dividing each sample's fluorescence value by the mean of the untreated control's fluorescence value. Red bars indicate the means of each group. Lower panels show representative images of CSP-stained sporozoites and their associated trails. Scale bars = 20 micrometers.
